# Improving Metagenomic Classification using Discriminative k-mers from Sequencing Data

**DOI:** 10.1101/2020.02.20.957308

**Authors:** D. Storato, M. Comin

## Abstract

The major problem when analyzing a metagenomic sample is to taxonomically annotate its reads in order to identify the species they contain. Most of the methods currently available focus on the classification of reads using a set of reference genomes and their k-mers. While in terms of precision these methods have reached percentages of correctness close to perfection, in terms of recall (the actual number of classified reads) the performances fall at around 50%. One of the reasons is the fact that the sequences in a sample can be very different from the corresponding reference genome, e.g. viral genomes are highly mutated. To address this issue, in this paper we study the problem of metagenomic reads classification by improving the reference k-mers library with novel discriminative k-mers from the input sequencing reads. We evaluated the performance in different conditions against several other tools and the results showed an improved F-measure, especially when close reference genomes are not available.

**Availability:** https://github.com/davide92/K2Mem.git

## 1 Introduction

Metagenomics is the study of the heterogeneous microbes samples (e.g. soil, water, human microbiome) directly extracted from the natural environment with the primary goal of determining the taxonomical identity of the microorganisms residing in the samples. It is an evolutionary revise, shifting focuses from the individual microbe study to a complex microbial community. The classical genomic-based approaches require the prior clone and culturing for the further investigation [10]. However, not all bacteria can be cultured. The advent of metagenomics succeeded to bypass this difficulty. Microbial communities can be analyzed and compared through the detection and quantification of the species they contain. In this paper we will focus on the detection of species in a sample using a set of reference genomes, e.g. bacteria and virus. The reference-based metagenomics classification methods can be broadly divided into two categories: (1) alignment-based methods, (2) sequence-composition-based methods, which are based on the nucleotide composition (e.g. *k*-mers usage). Traditionally, the first strategy was to use BLAST [1] to align each read with all sequences in GenBank. Later, faster methods have been deployed for this task, popular examples are MegaBlast [17] and Megan [7]. However, as the reference databases and the size of sequencing data sets have grown, alignment has become computationally infeasible, leading to the development of metagenomics classifiers that provide much faster results.

The fastest and most promising approaches belong to the composition-based one [9]. The basic principles can be summarized as follows: each genome of reference organisms is represented by its k-mers, and the associated taxonomic label of the organisms, then the reads are searched and classified throughout this k-mers database. For example, Kraken [15] constructs a data structure that is an augmented taxonomic tree in which a list of significant k-mers are associated to each node, leafs and internal nodes. Given a node on this taxonomic tree, its list of k-mers is considered representative for the taxonomic label and it will be used for the classification of metagenomic reads. Clark [12] uses a similar approach, building databases of species- or genus-level specific k-mers, and discarding any k-mers mapping to higher levels. The precision of these methods is as good as MegaBlast [17], nevertheless the processing speed is much faster [9]. Several other composition-based methods have been proposed over the years. In [5] the number of unassigned reads is decreased through reads overlap detection and species imputation. Centrifuge and Kraken 2 [8, 16] try to reduce the size of the k-mer database with the use respectively of FM-index and minimizers. The sensitivity can be improved by filtering uninformative k-mers [11,14] or by using spaced seeds instead of k-mers [3].

The major problem with these reference-based metagenomics classifiers is the fact that most bacteria found in environmental samples are unknown and cannot be cultured and separated in the laboratory [4]. As a consequence, the genomes of most microbes in an metegenomic sample are taxonomically distant from those present in existing reference databases. This fact is even more important in the case of viral genomes, where the mutation and recombination rate is very high and as a consequence the viral reference genomes are usually very different from the other viral genomes of the same species.

For these reasons, most of the reference-based metagenomics classification method do not perform well when the sample under examination contains strains that different from the genomes used as reference. Indeed, e.g. CLARK [12] and Kraken [15] report precisions above 95% on many datasets. On the other hand, in terms of recall, i.e. the percentage of reads actually classified, both Clark and Kraken usually show performances between 50% and 60%, and sometimes on real metagenomes just 20% of reads can be assigned to some taxa. In this paper we address this problem and we propose a metagenomics classification tool, named *K2Mem*, that is based, not only on a set of reference genomes, but also it uses discriminative k-mers from the input metagenomics reads in order to improve the classification. The basic idea is to add memory to a classification pipeline, so that previously analyzed reads can be of help for the classification.

## 2 Methods

To improve the metagenomics classification our idea is based on the following considerations. All reference-based metagenomics classification methods need to index a set of reference genomes. The construction of this reference database is based on a set of genomes, represented by its k-mers (a piece of genome with length k), and the taxonomic tree. For example, Kraken [15] constructs a data structure that is an augmented taxonomic tree in which a list of discriminative k-mers is associated to each node, leafs and internal nodes. Given a node on this taxonomic tree, its list of k-mers is considered representative for the taxonomic label and it is used for the classification of metagenomic reads. However, for a given genome only few of its k-mers will be considered discriminative. As a consequence, only the reads that contains these discriminative k-mers can be classified to this species.

Given a read with length *n*, each of its *n* − *k* + 1 k-mers have the first *k* − 1 bases in common with the previous k-mer, except the first k-mer. Furthermore, it is possible that reads belonging to the input sequencing data can have many k-mers in common.

As can be seen in Fig. 1, in this example we have three reads containing the same k-mer (in red) but only one is classified thanks to the presence in the read of a discriminative k-mer (in green), with a taxonomy ID associated, contained in the classifier’s database. The second read could not be classified because none of its k-mers are in the k-mer reference library, as there is a mutation (in bold) with respect to the reference genome. However, the k-mers of the first read, that are not present in the classifier’s database, can belong to the same species to which the read classified belongs to. With reference of the Fig. 1, if we associate to the shared k-mer (the red one) the taxonomy ID of the first read then, we can classify the other two reads. Thus, using the above considerations, one can try to extend the taxonomy ID of a classified read to all its k-mers.

**Fig. 1.**
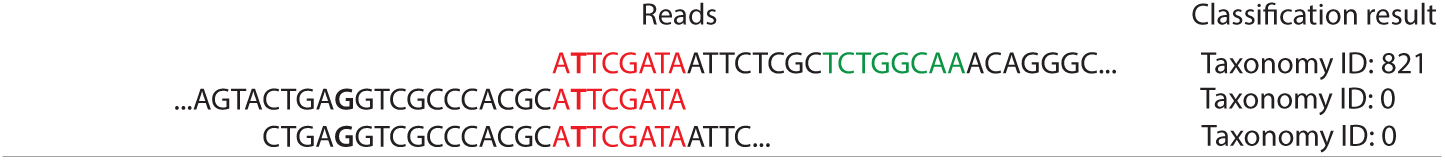
Example of a reference-based metagenomics classifier behavior. In red the k-mer in common between the reads, in green the k-mer associated to a species’ taxonomy ID (in this case 821) present in the classifier’s database, and in bold the mutations’ positions in the reads. A taxonomy ID of zero indicates that the classifier wasn’t able to classify the read.

The idea is to equip the classifier with memory from previous classifications, thus adding novel discrimintative k-mers found in the input sequencing data. To obtain this memory effect, one need to modify a given classifier with additional data structures and a new classification pipeline. Note that, this idea can be applied to any reference-based metagenomics classifiers that is based on a database of k-mers. In this paper we choose to use Kraken 2 [16], that was recently introduced, and that is reported to be the current state of the art. Before to describe our classification tool, for completeness here we give a brief introduction of Kraken 2 to better understand our contribution.

### 2.1 Background on Kraken 2

Kraken 2 is an improved version of the classifier Kraken [15] regarding memory usage and classification time. These improvements were obtained thanks to the use of the minimizers and a probabilistic compact data structure, instead of the k-mers and a sorted list used in Kraken.

At first, Kraken 2 needs to build a database starting from a set of reference genomes. To build the database Kraken 2 downloads from the NCBI the taxonomy and reference sequences libraries required. With the taxonomy data, Kraken 2 builds a tree with in each node a taxonomy ID. In each tree node a list of minimizer is stored that is useful for the classification. Precisely, for each minimizer (k=35, l=31) if it is unique to a reference sequence then, it is associated to the node with the sequence’s taxonomy ID. Instead, if the minimizer belongs to more than one reference sequence then, the minimizer is moved to the node with taxonomy ID equals to the Lowest Common Ancestor (LCA) of the two sequences it belongs. All the minimizer-taxonomy ID pairs are saved in a probabilistic Compact Hash Table (CHT) which allows a reduction of memory with respect to Kraken.

Once the database is built, Kraken 2 classifies the input reads in a more efficient manner than Kraken due to the smaller number of accesses to the CHT map. This is due to the fact that only distinct minimizers from the read trigger the research in the map. When the minimizer is queried in the compact table the node ID of the augmented tree is returned, and the counter in the node is increased. Once all the minimizers have been analyzed, the read is classified by choosing the deepest node from the root with the highest-weight path in the taxonomy tree.

### 2.2 K2Mem

Here we present K2Mem (Kraken 2 with Memory) a classifier based on Kraken 2. In order to implement the idea explained above, K2Mem needs to detect new discriminative minimizers, to store them in memory and to use these additional information in the classification of reads. The new classification pipeline of K2Mem discovers novel discriminative minimizers from the input sequencing data and it saves them in a map of additional minimizers.

The data structure used to store these additional minimizers is an unordered map that stores pairs composed of the novel discriminative minimizer, not present in the classifier’s database, and the taxonomy ID associated to the read that contains the minimizer. An unordered map is an associative container that contains key-value pairs with unique key. The choice of this structure is due to the fact that search, insertion and removal of elements have average constant-time complexity. Internally, the elements are not ordered in any particular order, but are organized in buckets. Which bucket an element is placed into depends entirely on the hash of its key. This allows a fast access to the single element, once the hash is computed, it refers to the exact bucket where the element has been inserted. The key and value are both 64 bit unsigned integer. This choice was made to keep the complete minimizers (l=31) on the map without loss of information due to the CHT hash function and to contains the taxonomy ID in case the number of taxonomy tree nodes increases in future.

K2Mem has two main steps, in the first phase all reads are processed and novel discriminative minimizers are discovered and stored in the additional minimizers map. In the second phase, the same input reads are re-classified using the Compact Hash Table and also the additional minimizers obtained in first phase.

An overview of the first phase, the discovery of novel discriminative minimizers, is reported in Figure 2. The population of the additional minimizers map works as follow: for each read, its minimizers (k=35, l=31) are computed one at a time and, for each of them, the Compact Hash Table (CHT) is queried (1). If it returns a taxonomy value equal to zero, then the additional map is queried if it is not empty (2). If the minimizer is not found in the additional minimizers map, this means that the minimizer is not in the Kraken 2’s database and no taxonomy ID has been assigned to it or is the first time the minimizer is found. In that case, the minimizer is added to a temporary list of not taxonomically assigned minimizers (3). Instead, if the CHT or the additional minimizers map query returns a taxonomy ID not equal to zero, then the taxonomy ID count is updated (4). Then, the read is classified (5), based on the highest-weight path in the taxonomy tree, and the resulting taxonomy ID is checked if it is at species level or below (6). If it is, then the minimizers in the unknown minimizers list are added to the additional minimizers map with key the minimizer and value the taxonomy ID obtained by the read classification. If the minimizer is already in the additional map the LCA of the input and stored taxonomy IDs value is saved. Instead, if the taxonomy ID obtained after the read classification is at a level above the species then, the minimizers are not added and the list is emptied. In the first phase this procedure is repeated for all the reads in the dataset. Then, once the population of the additional minimizers map is completed, in the second phase all reads are re-classified. In this phase the classification only uses the CHT and the additional minimizers map to classify the reads in the same dataset, but without to update the additional map content. With these additional minimizers, the dataset is processed to obtain a better classification, with reference to Figure 2 the classification pipeline follows the steps 1, 2, 4 and 5. When K2Mem ends, it generates the same output files as Kraken 2.

**Fig. 2.**
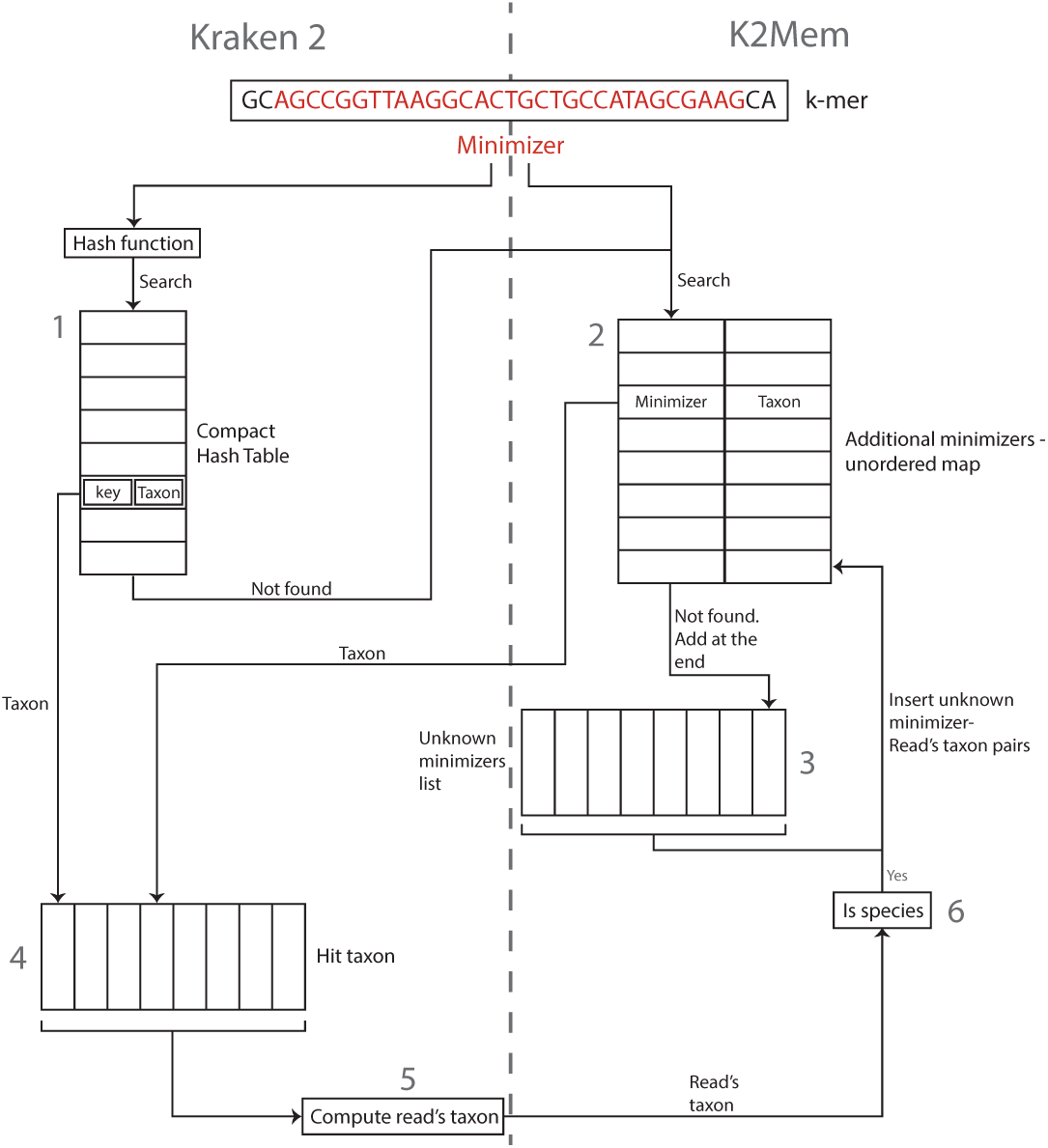
An overview of K2Mem and the discovery of additional minimizers.

## 3 Results

To analyze the performance of K2Mem we compare it with five of the most popular classifiers: Centrifuge [8], CLARK [12], Kraken [15], Kraken 2 [16], and KrakenUniq [2]. We test all these tools on several simulated datasets with different bacteria and viral genomes from NCBI. The experimental setup is described in next section.

### 3.1 Experimental setup

To assess the performance of K2Mem we follow the experimental setup of Kraken 2 [16], that is the strain exclusion experiment. This experiment consists in downloading the archaeal, bacteria, and viral reference genomes and the taxonomy from NCBI. From this set of reference genomes 50 are removed (40 bacteria and 10 viruses), called the origin strains, that are used for testing. All the remaining reference genomes are used to build the classifiers’ databases, one of each tool in this study. This setup tries to mimic a realistic scenario in which a reference genome is available, but the sequenced genome is a different strain from the same species.

With the origin strains chosen we build 10 datasets using Mason 2 [6] to simulate 100 bps paired-end Illumina sequence data. Of these datasets, 7 are built by varying the number of reads, from 50k to 100M, using the default Illumina error profile. These datasets are used to test the impact of sequencing coverage on the performance of the tools under examination. We also constructed other 3 datasets, all with the same number of reads 100M, but with different mutation rates from the original strains: 2%, 5%, and 10%. With these datasets we evaluate another scenario in which a close reference genome is not available.

To compare K2Mem with the other tools we use the same evaluation metrics as in [14, 16]; precisely we use Sensitivity, Positive Predicted Value (PPV), F-measure, and Pearson Correlation Coefficient (PCC) of the abundance profile.

### 3.2 Performance Evaluation

In this section we analyze the performance results at genus level of K2Mem with respect to the other classifiers. All tools are used with the default parameters and run in multithreading using 16 threads. Their databases are built using the same set of reference genomes obtained from the strain exclusion experiment. The obtained results are reported below in different figures to better understand the performance and the impact of the different configurations.

In Figure 3 and 4 are shown the full results obtained with the 100M reads dataset for bacteria and viruses respectively. We analyze the 100M dataset as the behavior of K2Mem with the other datasets is similar. As it can be seen in Figure 3, with bacteria K2Mem obtains an F-measure improvement of at least 0.5 percentage points (pps) respect to the closest competitor, Kraken 2, and the other classifiers. This improvement is due to an increase of sensitivity of at least 2 pps despite a worsening of the PPV of about 1 pps with respect to the other tools. Moreover, K2Mem obtains the best PCC value with an improvement of at least 0.1 pps respect to Centrifuge and Kraken 2.

**Fig. 3.**
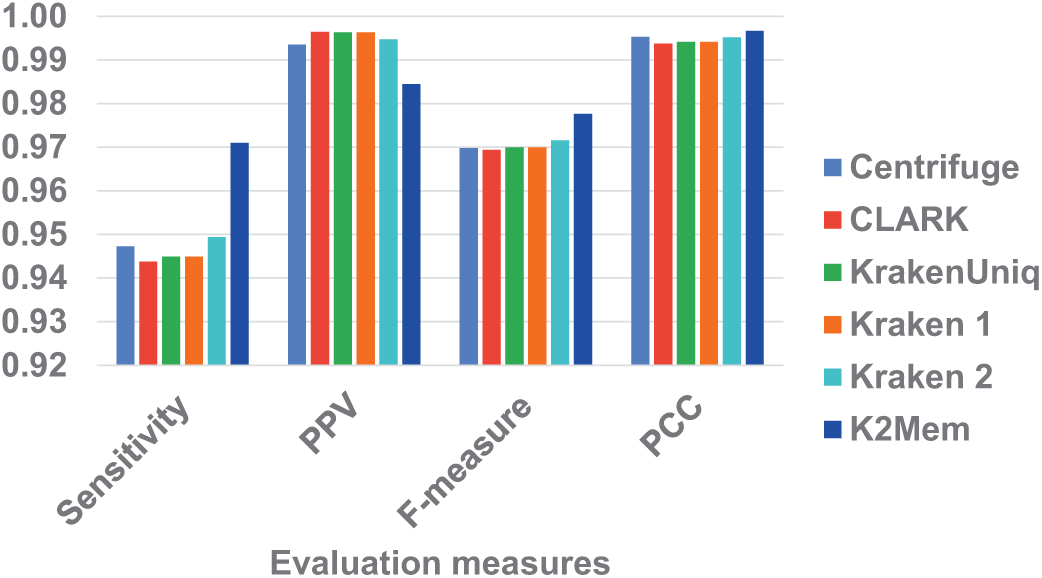
Bacteria evaluation at genus level on the 100M reads dataset.

**Fig. 4.**
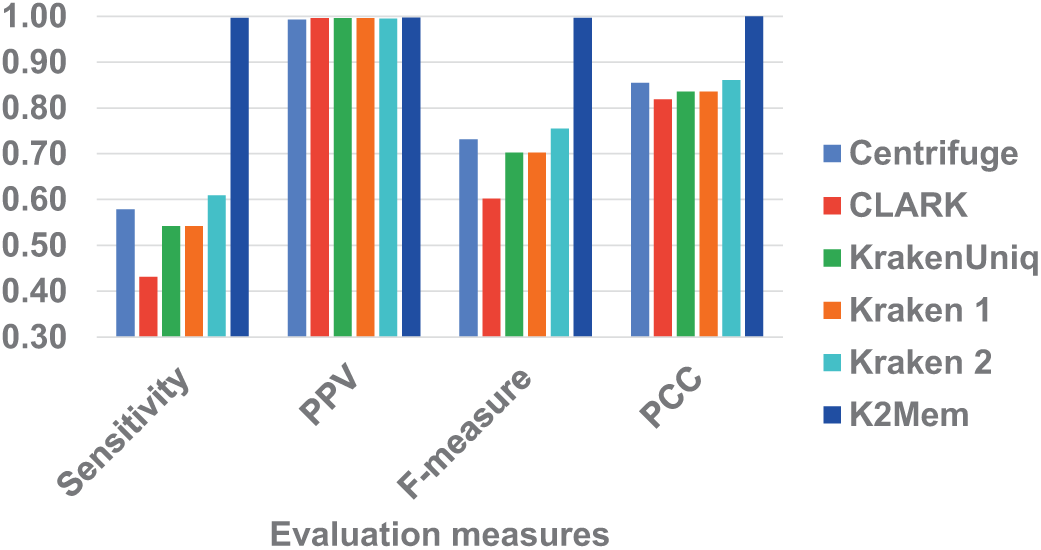
Viruses evaluation at genus level on the 100M reads dataset.

On the viral dataset, as it can be observed in Figure 4, K2Mem obtains an F-measure improvement of almost 40 pps with respect to Kraken 2 and the other tools. This improvement is due to an increase of sensitivity, whereas the PPV of all tools is to close to 1. Moreover, K2Mem shows the best PCC with an improvement of about 13 pps respect to Centrifuge and Kraken 2.

In summary, with the results reported above we can observe that thanks to the additional information provided by the new discriminative minimizers K2Mem obtains a moderate improvement in bacteria’s classification and a significant improvement in viruses’ classification. Similar results are observed also for species level classification (data not shown for space limitation).

In Figure 5 and 6 are shown the F-measure values obtained varying the number reads in the dataset for bacteria and viruses respectively. For bacteria, as it can be seen in Figure 5, K2Mem gets better F-measure values than the other tools as the number of reads increases; obtaining improvements up to almost 1 pps. This improvement is given mainly from the increase of sensitivity.

**Fig. 5.**
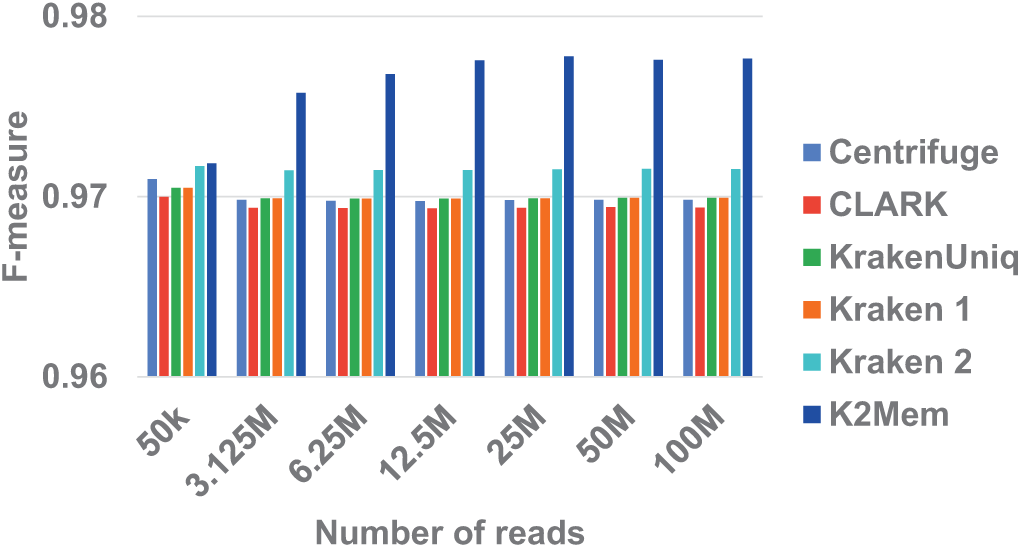
F-measure values for bacteria at genus level varying the number of reads.

**Fig. 6.**
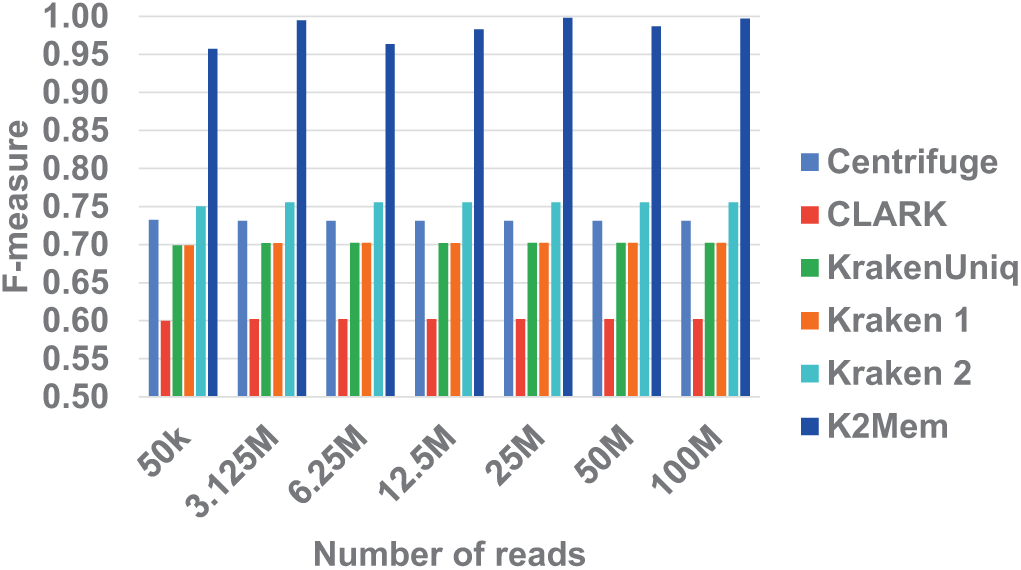
F-measure values for viruses at genus level varying the number of reads.

For viruses, as it can be seen in Figure 6, K2Mem achieves greater F-measure improvement than bacteria, always thanks to an increase of the sensitivity. It is interesting to note that the performance of all other tools are independent on the size of the dataset. However, we can observe that for K2Mem the greater the amount of data the better the classification results, due to a greater possibility of finding new discriminative minimizers.

In Figure 7 and 8 are shown the F-measure values obtained for the 100M reads dataset while varying the mutation rate, for bacteria and viruses respectively. The mutation rate with respect to the origin strains rages between 0.4% to 10%.

**Fig. 7.**
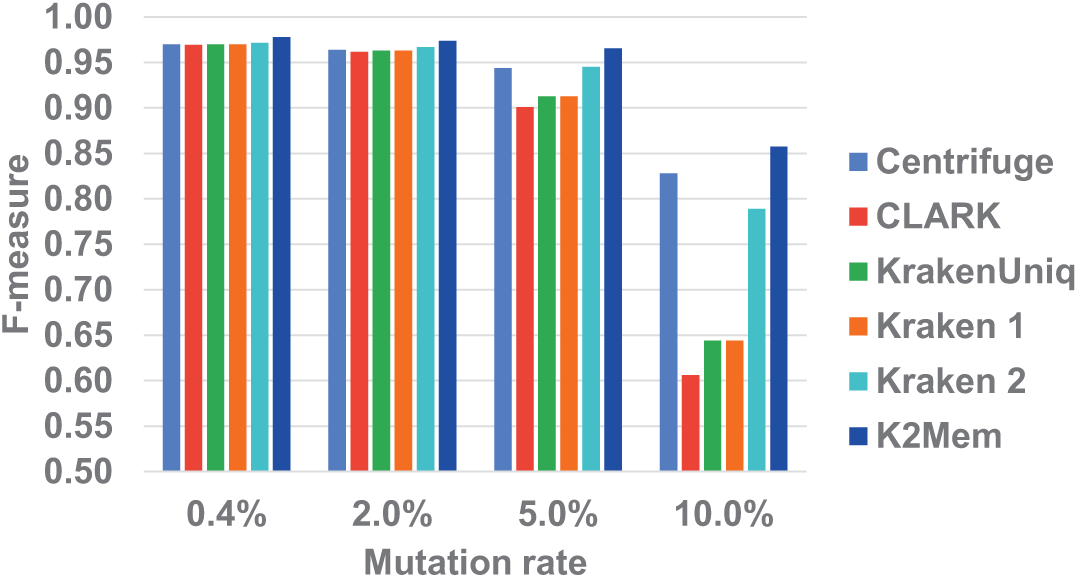
F-measure values for bacteria at genus level varying the mutation rate.

**Fig. 8.**
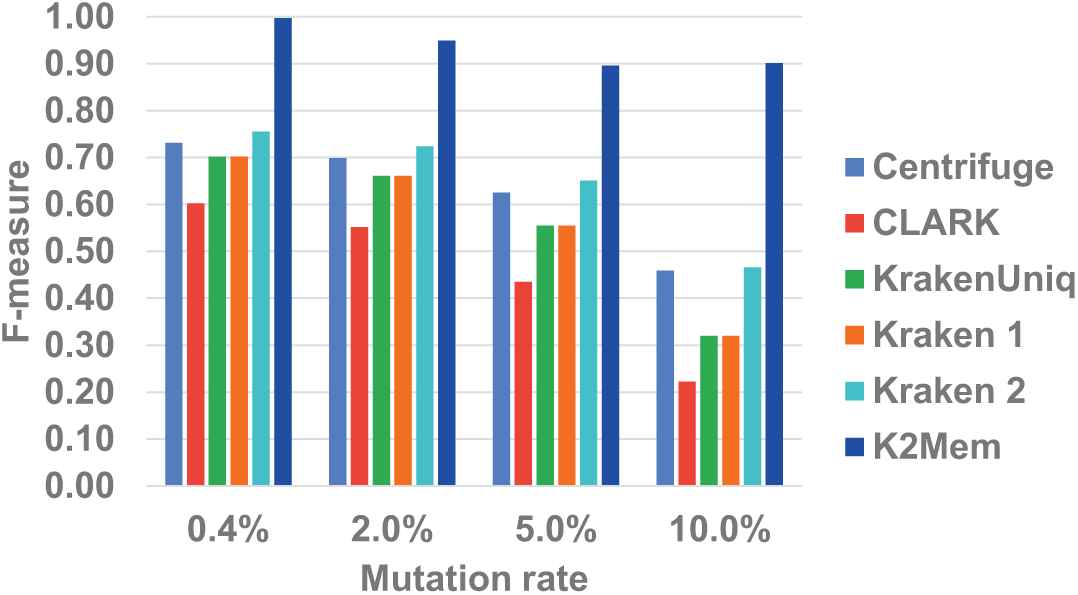
F-measure values for viruses at genus level varying the mutation rate.

As it can be seen in Figure 7, the first observation is that the performance of all tools decreases as the mutation rate increases. This is expected, because the reference genomes are no longer similar to the genomes in the sample data. However, K2Mem obtains the best F-measure values in all the cases. Moreover, the F-measure improvement increases up to 2.5 pps w.r.t. Centrifuge and 7 pps with Kraken 2, as the mutation rate increases.

With viruses, as reported in Figure 8, K2Mem has the same behavior described for the bacteria, but with a bigger performance gap up to 45 pps with respect to Kraken 2. From the results above, the increase of the mutation rate leads to worse performance for all tools as expected. However, K2Mem is the classifier that has suffered less from the presence of mutations in the sample.

### 3.3 Execution time and memory usage

The execution time and memory usage of each tool during the datasets classification are showed in Figure 9. For this analysis the execution time and memory usage values are reported for the largest dataset with 100M reads.

**Fig. 9.**
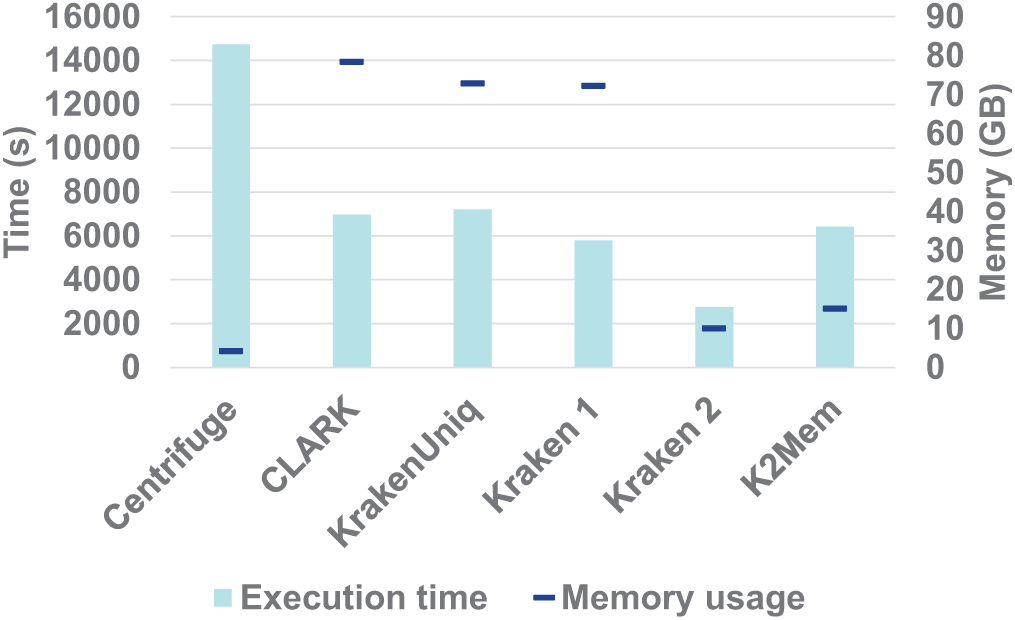
Execution time and memory usage for a dataset with 100M reads.

The execution time of K2Mem, as expected, it is larger than Kraken 2 due to the new discriminative minimizers search phase. However, the classification time is in line with the other tools. As for the memory usage K2Mem needs more memory than Kraken 2, due to the new map. However, K2Mem requires only 15 GB of RAM in line with the best tools.

## 4 Conclusions

We have presented K2Mem, a classifier based on Kraken 2 with a new classification pipeline and an additional map to store new discriminative k-mers from the input sequencing reads. The experimental results has demonstrated that K2Mem obtains higher values of F-measure, mainly by an improved sensitivity, and PCC with respect to the most popular classifiers thanks to the greater number of reference k-mers available during the classification. We showed that the performance improvement of K2Mem increases as the size of the input sequencing data grows, or when reads originate from strains that are genetically distinct from those in the reference database. As possible future developments it could be interesting to increase the PPV, e.g. using unique k-mers, to speedup the classification algorithm with a better implementation and to test other data structures, e.g. counting quotient filter [13], to decrease the memory requirements.

## References

1. Altschul, S.F., Gish, W., Miller, W., Myers, E.W., Lipman, D.J.: Basic local alignment search tool. Journal of Molecular Biology 215(3), 403–410 (1990)

2. Breitwieser, F., Baker, D., Salzberg, S.L.: Krakenuniq: confident and fast metagenomics classification using unique k-mer counts. Genome biology 19(1), 198 (2018)

3. Břinda, K., Sykulski, M., Kucherov, G.: Spaced seeds improve k-mer-based metagenomic classification. Bioinformatics 31(22), 3584 (2015). https://doi.org/10.1093/bioinformatics/btv419

4. Eisen, J.A.: Environmental shotgun sequencing: its potential and challenges for studying the hidden world of microbes. PLoS Biol. 5 (2007)

5. Girotto, S., Comin, M., Pizzi, C.: Higher recall in metagenomic sequence classification exploiting overlapping reads. BMC Genomics 18(10), 917 (Dec 2017). https://doi.org/10.1186/s12864-017-4273-6

6. Holtgrewe, M.: Mason: a read simulator for second generation sequencing data (2010)

7. Huson, D.H., Auch, A.F., Qi, J., Schuster, S.C.: Megan analysis of metagenomic data. Genome Res. 17 (2007)

8. Kim, D., Song, L., Breitwieser, F., Salzberg, S.: Centrifuge: Rapid and sensitive classification of metagenomic sequences. Genome Research 26, gr.210641.116 (10 2016). https://doi.org/10.1101/gr.210641.116

9. Lindgreen, S., Adair, K., Gardner, P.: An evaluation of the accuracy and speed of metagenome analysis tools. Cold Spring Harbor Laboratory Press (2015)

10. Mande, S.S., Mohammed, M.H., Ghosh, T.S.: Classification of metagenomic sequences: methods and challenges. Briefings in Bioinformatics 13(6), 669–681 (Nov 2012). https://doi.org/10.1093/bib/bbs054

11. Marchiori, D., Comin, M.: Skraken: Fast and sensitive classification of short metagenomic reads based on filtering uninformative k-mers. In: BIOINFORMATICS 2017 - 8th International Conference on Bioinformatics Models, Methods and Algorithms, Proceedings; Part of 10th International Joint Conference on Biomedical Engineering Systems and Technologies, BIOSTEC 2017. vol. 3, pp. 59–67 (2017)

12. Ounit, R., Wanamaker, S., Close, T.J., Lonardi, S.: Clark: fast and accurate classification of metagenomic and genomic sequences using discriminative k-mers. BMC Genomics 16(1), 1–13 (2015). https://doi.org/10.1186/s12864-015-1419-2

13. Pandey, P., Bender, M.A., Johnson, R., Patro, R.: A general-purpose counting filter: Making every bit count. In: Proceedings of the 2017 ACM International Conference on Management of Data. pp. 775–787. ACM (2017)

14. Qian, J., Marchiori, D., Comin, M.: Fast and sensitive classification of short metagenomic reads with skraken. In: Peixoto, N., Silveira, M., Ali, H.H., Maciel, C., van den Broek, E.L. (eds.) Biomedical Engineering Systems and Technologies. pp. 212–226. Springer International Publishing, Cham (2018)

15. Wood, D., Salzberg, S.: Kraken: ultrafast metagenomic sequence classification using exact alignments. Genome Biol. 15 (2014). https://doi.org/10.1186/gb-2014-15-3-r46

16. Wood, D.E., Lu, J., Langmead, B.: Improved metagenomic analysis with kraken 2. Genome biology 20(1), 257 (2019)

17. Zhang, Z., Schwartz, S., Wagner, L., Miller, W.: A greedy algorithm for aligning dna sequences. Journal of Computational Biology 7 (1-2), 203–214 (2004)

